# Patch size distribution and network connectivity affect species invasion dynamics in dendritic networks

**DOI:** 10.1101/2020.01.13.903070

**Authors:** Kathrin Holenstein, Eric Harvey, Florian Altermatt

## Abstract

Biological invasions are globally affecting ecosystems, causing local species loss and altering ecosystem functioning. Understanding the success and unfolding of such biological invasions is thus of high priority. Both local properties and the spatial network structure have been shown to be determinants of invasion success, and the identification of spatial invasion hubs directly promoting invasion dynamics is gaining attention. Spatial dynamics, however, could also indirectly alter invasion success by shaping local community structure: in many ecosystems, such as riverine networks, regional properties such as patch size distribution are known drivers of local community structures, which themselves may affect the establishment success of invading species. Using microcosm experiments in dendritic networks, we disentangled how patch size distribution and dispersal along specific network topologies shaped local communities, and, subsequently, affected the establishment success of invading species. We find that inherent patch size distributions shaped composition and diversity of local communities, and, subsequently, modulated invasion success. Specifically, the relationship between local diversity and invasion success changed across an increasing patch size gradient from a negative to a positive correlation, while overall increasing patch size reduced invasion success. Connectivity did not have a direct effect on invasion success but indirectly affected invasions by shaping diversity patterns in the whole network. Our results emphasize the relevance of indirect, landscape-level effects on species invasions, which need to be considered in the management of spatial habitat networks.

## Introduction

Biodiversity is rapidly changing worldwide and declines are predicted to continue for the coming decades (Chapin et al. 2000, Sala et al. 2000, Pereira et al. 2010). One major factor contributing to this decline are biological invasions, often occurring as a bi-product of increased human mobility, global trade, and changes in habitat or ecosystem properties (Sala et al. 2000, Mooney and Cleland 2001). Species invading new ecosystems can partially or completely out-compete, predate, or displace native species, with subsequent impacts on ecosystem functioning (Clavero and García-Berthou 2005, Pereira et al. 2010), calling for a mechanistic understanding of species invasion success and the ecological drivers of it.

Invasion success has been linked to local properties of the communities and ecosystems (Stachowicz et al. 1999, Levine et al. 2004, Mächler and Altermatt 2012, Kempel et al. 2013), but also to regional properties and spatial landscape structures (Deckers et al. 2005, Mari et al. 2014, Altermatt and Fronhofer 2018). The former is relatively straightforward: local conditions related to resource availability and the functional niche of each resident species will directly influence the establishment success of an invading species. As such, biodiversity and other properties of the community have been proposed as important local modulators of species invasion success (Levine and D’Antonio 1999, Stachowicz et al. 1999, Levine et al. 2004). Overall, most studies indicate a positive effect of species diversity in reducing species invasions (Elton 1933, Tilman 1997, Wardle 2001, Kennedy et al. 2002), mostly because a diverse community is assumed to already have filled the available niches with resident species efficiently using the available resources, such that it is more difficult for an invader to establish and persist.

The latter, effects of regional connectivity and the spatial landscape structure, is more complex. Firstly, spatial dynamics directly modulate how invading species get to a specific location, and thereby affect invasion dynamics. In this context, the spatial unfolding of invasion fronts (Giometto et al. 2014, Carraro et al. 2018) as well as the significance of specific hubs in spatial networks on invasion success has been receiving increasing attention (e.g., Morel-Journel et al. 2019). Secondly, however, the physical configuration of the landscape and the spatial dynamics it generates will also modulate invasion success indirectly by influencing local community structure and dynamics such as diversity or identity of species found locally (MacArthur and Wilson 1963, Holoyak et al. 2005, Holland and Hastings 2008, Pillai et al. 2011, McIntosh et al. 2018). As virtually all species live in a spatial landscape of connected habitats, this spatial perspective is highly relevant. These intricate effects of spatial dynamics on invasion success are expected in all spatially structured landscapes, but may be especially pronounced in spatial networks with a complex but also non-random spatial structure (Rodriguez-Iturbe and Rinaldo 1997).

A key candidate for such landscapes are dendritic riverine networks, which are most strongly affected by invasive species (Leuven et al. 2009, Reid et al. 2019). They follow well described network structures and properties that are globally highly conserved (Rodriguez-Iturbe and Rinaldo 1997, Altermatt 2013). It is well known that dispersal and metacommunity dynamics are essential to properly describe the spatial linkage between communities in such riverine networks (Altermatt 2013, Tonkin et al. 2018). Also, connectivity and network structure are known to have a central role in shaping local population dynamics and species composition (Corre et al. 2015, Layeghifard et al. 2015, Altermatt and Fronhofer 2018). Specifically, riverine landscapes possess typical and invariant characteristics leading to higher species diversity at confluences of branches and in lower reaches of the stream (Fernandes et al. 2004, Muneepeerakul et al. 2007, Rodriguez-Iturbe et al. 2009). Experiments in dendritic landscapes have shown that those diversity patterns are primarily shaped by the connectivity of habitats and the directionality of dispersal between habitats (Carrara et al. 2014, Seymour et al. 2015), but can also be modulated by disturbances (Harvey et al. 2018).

Consequently, it has been speculated that predicting invasion dynamics requires a better understanding of regional influences related to landscape network structure (Mari et al. 2014). Previous work has thus independently studied the relationships between habitat size and diversity, connectivity and diversity, and the effect of species diversity on invasion success (Kennedy et al. 2002, Drakare et al. 2006, Carrara et al. 2012, Meier and Hofer 2016). However, little is known about the interactive effects of habitat connectivity, habitat size, and species diversity on invasion success.

In this study, we experimentally tested how well-conserved habitat size distribution and connectivity in dendritic networks are shaping local community composition, and, subsequently, affecting the resistance of these communities to species invasions. We conducted microcosm experiments with protist species, in which we first let the local communities assemble across replicated dendritic networks with varying patch size distribution and respective isolated controls. Subsequently, we stopped dispersal in the dendritic network and introduced invaders into each of the local communities. This allowed us to address the invasibility of each community, and to disentangle effects of local community properties *per se* and the regional dynamics influencing resident community assembly *prior* to invasion. We assessed short- and long-term invasion success. The invasion success was assessed for each local community individually, thereby disentangling effects of local properties and the imprint of spatial network position. We investigated the effect of patch connectivity and patch size on the establishing of the invaders using presence and absence data of invaders. We hypothesized that the invasion success is lower in large patches. Those larger patches are expected to support higher species diversity, with assumed higher resource complementarity, higher proportion of used resources and therefore lower availability of resources for invaders (Wardle 2001). On the contrary, the invasion success was expected to be higher in small patches with low resource complementarity and low species diversity, a low proportion of used resources, and higher resource availability for invaders. We also assumed the imprint of spatial connectivity on local communities to subsequently reduce invasion success, because past dispersal dynamics (compared to the isolated controls) should have made communities more resistant to invading species.

## Methods

We studied community assembly and subsequent invasion dynamics in experimental dendritic networks (Fig. 1). We used five independent realizations of dendritic network landscapes (A to E), with 36 patches each (Fig. S1 Appendix). The structure of these networks followed optimal channel structures with patch size scaling distributions observed in real river systems, which have already been used in previous experiments (Carrara et al. 2014). For logistic reasons, patch sizes were binned in four categories (3 mL, 5.2 mL, 9 mL, 18 mL), and as a control we had 10 isolated replicates (isolated “control”) for each of these patch sizes (in total 40 control patches; Fig. 1).

**Figure 1:**
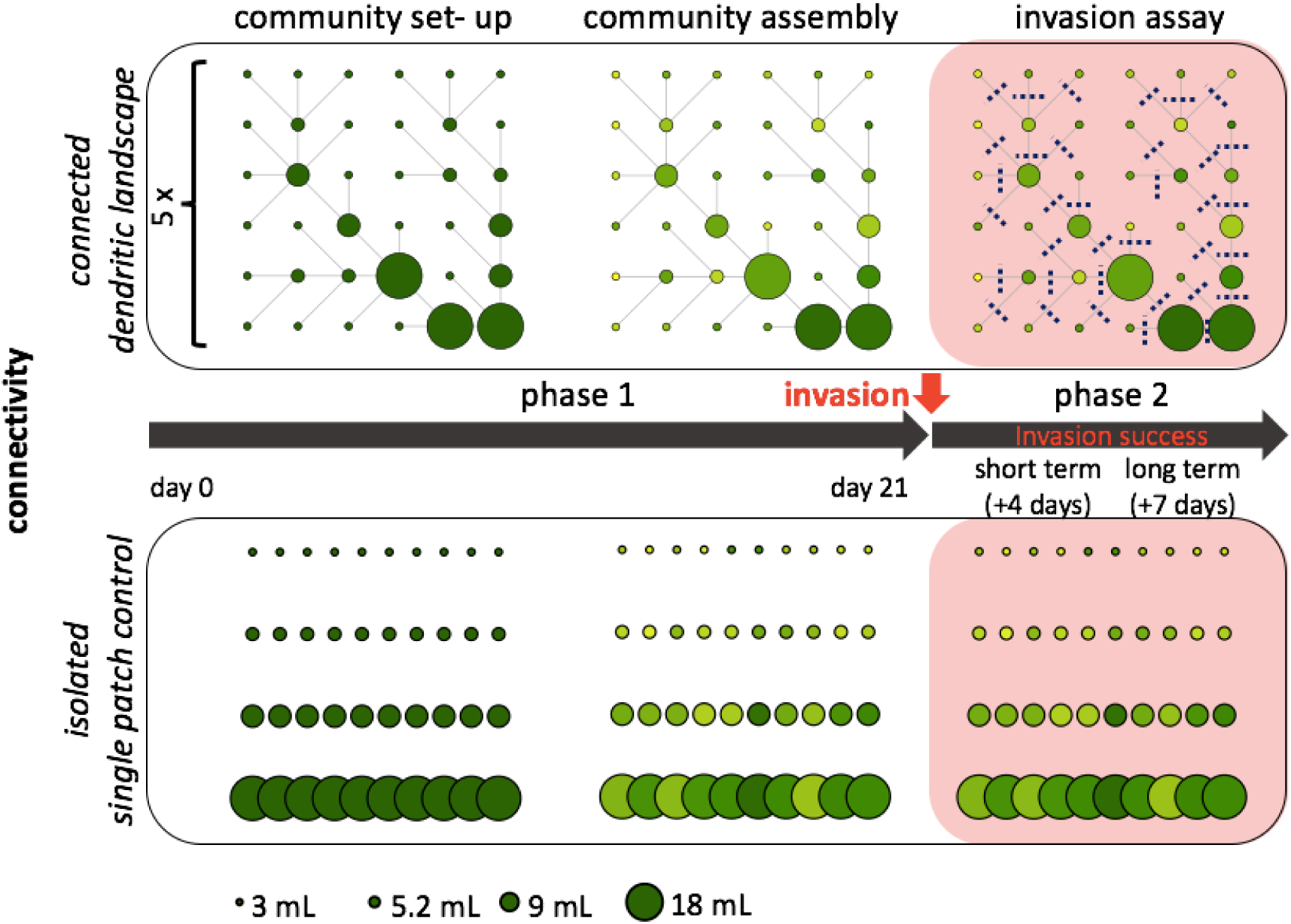
Experimental setup. Five dendritic landscapes with 36 connected patches following a realistic patch size scaling structure as well as 40 single patch controls were each inoculated with the same community of six protist species. Subsequent community structure and biodiversity was shaped by dispersal and species interactions over a period of 21 days (single patch controls without dispersal). Thereafter, dispersal was halted, and the species *Spirostomum* sp. was added to each patch, and its invasion (establishment) success was assessed at two subsequent time points.

We introduced six ‘resident’ protist species into each patch, monitored subsequent community assembly shaped by dispersal, species interactions, and ecological drift (phase 1; Fig. 1). We then halted dispersal, and assessed the invasibility potential of each of these communities, by introducing an invading species to each of these communities after three weeks of community assembly (phase 2). We investigated invasion success by measuring presence or absence of the invader on day four (∼8 generations, termed short-term invasion success) and day seven post-invasion (∼14 generations, termed long-term invasion success). Importantly, this design allowed us to separate the effect of spatial dynamics on community assembly (phase 1) from the subsequent test of invasion potential (phase 2), which was done for each of the patches independently.

### Community set-up and community assembly

We used six freshwater protist and rotifer species for the community assembly: *Loxocephalus* sp. (Lox), *Tetrahymena* sp. (Tet), *Colpidium striatum* (Col), *Dexiostoma campylum* (Dex), *Paramecium caudatum* (Pca) and *Cephalodella* sp. (Rot, a rotifer, in the following referred to as protist). These protists were kept in a nutrient medium inoculated with the bacteria *Serratina fonticola, Brevibacillus brevis*, and *Bacillus subtilis* as a food source. All species had been kept as monocultures and kept under sterile conditions following Altermatt et al. (2015).

At the onset of the experiment, we mixed the six protist species monocultures with a focal density of 33.3 individuals mL^−1^ per species (community-mix). From this community-mix we distributed 3 mL to all patches, such that it contained on average 100 individuals of each of the six species Lox, Tet, Col, Dex, Pca, and Rot. Starting at these densities, well-below carrying capacity, allowed us to look at species assembly dynamics. We toped up the volume in the respective patches with protist medium to reach the final focal volumes (3 mL, 5.2 mL, 9 mL, 18 mL).

We then applied a dispersal treatment twice a week (in total 5 times during 21 days). We applied undirected nearest neighbor dispersal along the network structure, using a method with mirror landscapes developed by Carrara et al. (2012) to avoid long-tailed dispersal. This dispersal treatment was expected to shape community assembly by counteracting local extinctions due to species interactions or ecological drift. At each dispersal step, patches were well-mixed and then a volume of 200 µL was transferred into each neighboring patch using the mirror landscape (Fig. S2 Appendix). In the isolated single patch controls no dispersal was conducted, but a volume of 200 µL was pipetted out and back into the same patch to control for the handling effect.

### Invasion assay

Based on results from a preliminary experiment, *Spirostomum* sp. was chosen as the invader to these communities. This species was chosen as it is, also in comparison with the other species, relatively large (body length ∼850 µm, for traits of all species see also Carrara et al. 2012), and thus can be tracked easily in the communities. 21 days after the onset of the experiment, we stopped all dispersal, and added approximately 10 *Spirostomum* sp. individuals in a volume of 200 µL to each patch. We intentionally added this invading species at an initially low number to match natural scenarios of species invasions. Four days and seven days after this invasion, presence or absence of the invader was assessed (short-term invasion success and long-term invasion success). To do so, we screened the invader’s presence in a subsample of 175 µL under a stereo microscope (Leica M205 C, Leica Microsystems GmbH, Wetzlar DE). When *Spirostomum* sp. was not present in the subsample screened for the long-term invasion success, we additionally screened a total volume of 1.5 mL.

In parallel, we used a video-based monitoring method to assess community structure and diversity of resident communities. We recorded and analyzed the community in each patch of the connected dendritic landscape and isolated single patch controls at three different time points: immediately preceding the invasion, four days after the invasion, and seven days after the invasion. The videos for community analysis were taken from 175 µL subsamples, of which a total volume of 34.4 µL was recorded for 5 seconds (25 frames per second, 16x fold magnification, full light) by a digital Orca Flash 4.0 camera (C11440-22CU, Hamamatsu Photonics, Japan). We closely followed the method and the R package BEMOVI developed and used by Pennekamp, Schtickzelle, and Petchey (2015) and Pennekamp et al. (2017). The settings for BEMOVI script were the following: Pixel size of 4.05 µm, difference lag of 10 frames, thresholds of 10 to 255 difference of pixel intensity, min particle size 5 pixels, max particle size 1000 pixels, link range 3 frames, displacement 16 pixels, detection frequency of 0.1 seconds, median step length of 3 pixels.

To identify species, we used a random forest algorithm. This algorithm is based on decision trees using binary thresholds to divide the observations into the most possible class at the end node (Pennekamp et al. 2017). The information about morphological and movement features for classification were given from BEMOVI (Pennekamp et al. 2017).

### Statistical analyses

We calculated Shannon diversity of the community in every patch using the R-package vegan (version 2.4-3, Oksanen et al. 2017). The degree connectivity of each patch (i.e., number of connecting nodes) and distance to outlet was calculated directly from the respective network adjacency matrices.

We used the R-package lme4 (version 1.1-13, Bates et al. 2015), to analyze factors explaining observed patterns of diversity (Shannon diversity) in the landscapes as well as subsequent invasion success, the latter based on presence and absence data. We ran linear mixed effect models (lmer) to analyse Shannon diversity, including the explanatory variables patch size, connectivity, and distance to outlet. We ran generalized linear mixed-effects models (glmer) with a binomial distribution to analyse invasion success, including the explanatory variables patch size, connectivity, distance to outlet, and Shannon diversity immediately preceding invasion. Landscape identity was added as a random factor. We did the model selection based on the AICc and its weights using the R-package MuMIn (version 1.43.6, Barton 2019), and looked at the relative variable importance that sums AICc weights over all models and explanatory variables. Based on the best-fitting model and its variance-covariance matrix, we did model predictions for invasion success as a function of diversity before invasion using the R-package mvtnorm (version 1.0-11, Genz et al. 2017) generating random numbers for the predictions. To compare the proportion of invasion success between communities in different patch sizes of the landscape and the control, we used a chi-square test. All statistical analyses were done in R (version 3.3.3, R Development Core Team. 2017).

## Results

Dispersal in the dendritic landscapes affected community assembly and resulted in a characteristic diversity distribution: Shannon diversity index (S_H_) immediately preceding the invasion varied between 0.32 to 1.42, with smallest values found in small and/or more isolated patches (Fig. 2), while highest values were found in the largest patches. Overall, in dendritic landscape, mean S_H_ immediately preceding invasion increased steadily from small (3 mL) to large (18 mL) patches (mean S_H_ with increasing patch size: 0.70, 0.74, 0.96, 1.09, Figure S3 Appendix). This is also seen in the model selection, where patch size is the single most important factor explaining diversity, followed by connectivity and distance to outlet, which were also retained in all models as important factors explaining diversity (Table 1). Diversity in single patch controls was also significantly increasing with patch size, (Figure S3 Appendix) but was on average significantly lower (p<0.03; all statistical details in Table S1 Appendix) in the isolated control compared to the diversity in the dendritic network.

**Table 1:** Importance table for the analysis of Shannon diversity preceding invasion. The sum of AICc weights over all the models is given for all variables, ordered in decreasing importance. Also, the number of models retaining each variable is given. Variables assessed were patch size, connectivity, distance to outlet, and all their interaction terms (:).

| Variable | Importance | N models |
| --- | --- | --- |
| Patch size | 1 | 14 |
| Connectivity | 0.01 | 14 |
| Distance to outlet | 0.01 | 14 |
| Distance to outlet : Patch size | <0.01 | 6 |
| Connectivity : Distance to outlet | <0.01 | 6 |
| Connectivity: Distance to outlet : Patch size | <0.01 | 1 |

**Figure 2:**
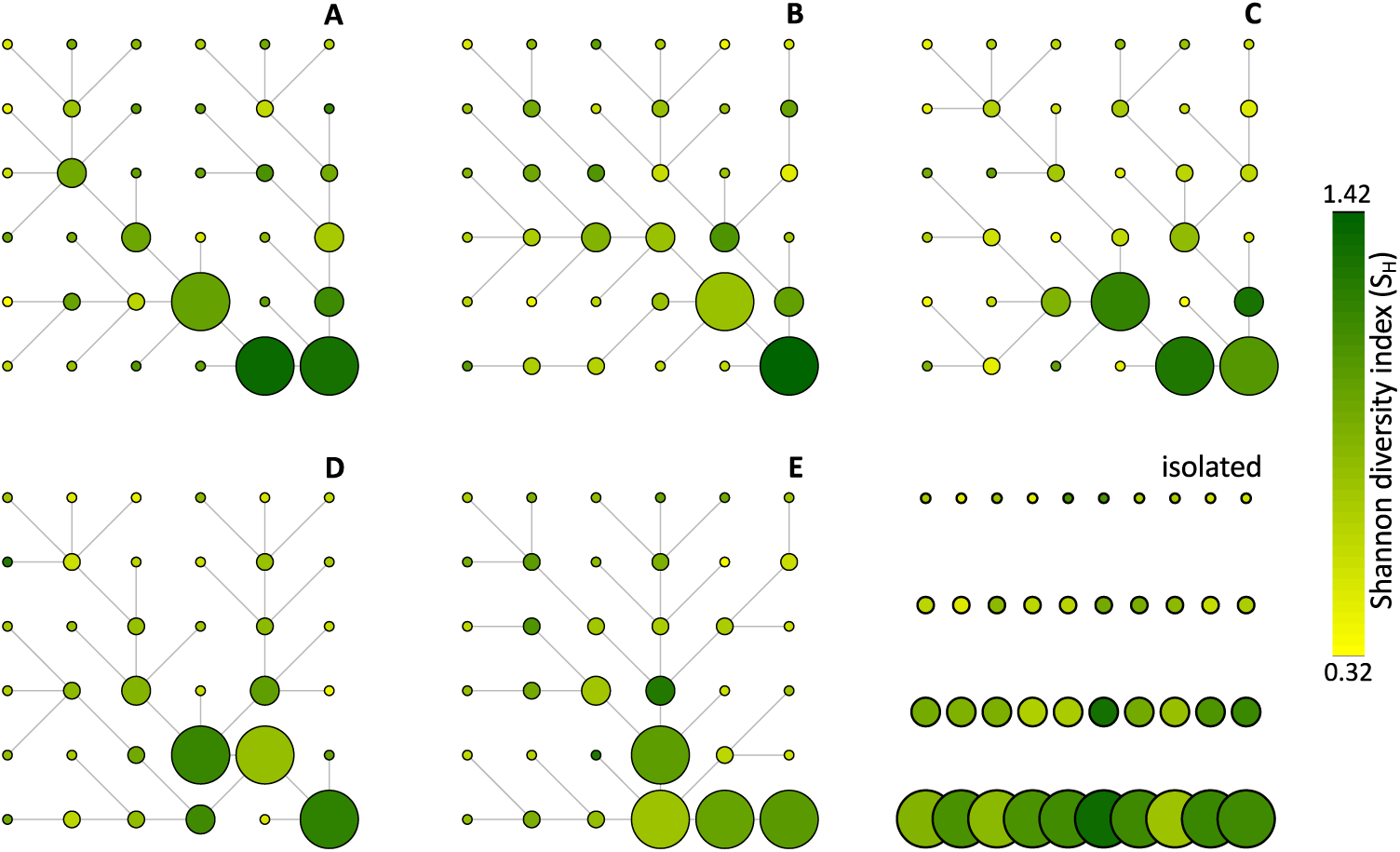
Shannon diversity (S_H_) of the resident communities immediately preceding the invasion of *Spirostomum* sp. across the dendritic landscapes (A to E) and the isolated single patch controls.

We then analyzed invasion success of Spi in the landscapes (Fig. 3). The short-term invasion success of Spi in the dendritic landscapes was highest in the small patches (3 mL) and lowest in the large patches (18 mL). The mean proportion of successful invasions was 9.5 times higher in the small 3 mL patches (0.57) compared to the 18 mL patches (0.06). Based on the sum of weights of AICc, patch size was the most important variable explaining the short-term invasion success of Spi. Shannon-diversity (preceding the invasion), connectivity (here see also Fig. S4 Appendix) and distance to the outlet were similarly important as explanatory variables explaining short-term invasion success, and were all retained in all models (Table 2). The results for long-term invasion success were highly congruent (Table 3), with the same variables, and almost same order of importance emerging (Fig. S5 Appendix).

**Table 2:** Importance table for the analysis of short-term invasion success. The sum of AICc weights over all the models is given for all variables, ordered in decreasing importance. Also, the number of models retaining each variable is given. Variables assessed were patch size, Shannon diversity index (SH) immediately preceding invasion, connectivity, distance to outlet, and all their interaction terms (:).

| Variable | Importance | N models |
| --- | --- | --- |
| Patch size | 1 | 148 |
| SH | 0.74 | 148 |
| SH : Patch size | 0.61 | 84 |
| Connectivity | 0.52 | 148 |
| Distance to outlet | 0.47 | 148 |
| Connectivity : Patch size | 0.17 | 84 |
| Distance to outlet : Patch size | 0.16 | 84 |
| Connectivity : SH | 0.12 | 84 |
| Distance to outlet : SH | 0.1 | 84 |
| Connectivity : Distance to outlet | 0.1 | 84 |
| Connectivity : SH : Patch size | <0.01 | 20 |
| Distance to outlet : SH : Patch size | <0.01 | 20 |
| Connectivity : Distance to outlet : Patch size | <0.01 | 20 |
| Connectivity : Distance to outlet : SH | <0.01 | 20 |
| Connectivity : Distance to outlet : SH : Patch size | <0.01 | 1 |

**Table 3:** Importance table for the analysis of long-term invasion success. The sum of AICc weights over all the models is given for all variables, ordered in decreasing importance. Also, the number of models retaining each variable is given. Variables assessed were patch size, Shannon diversity index (SH) immediately preceding invasion, connectivity, distance to outlet, and all their interaction terms (:).

| **Variable** | **Importance** | **N models** |
| --- | --- | --- |
| Patch size | 0.9 | 148 |
| Distance to outlet | 0.69 | 148 |
| SH | 0.58 | 148 |
| Connectivity | 0.48 | 148 |
| Distance to outlet : SH | 0.17 | 84 |
| Distance to outlet : Patch size | 0.16 | 84 |
| SH : Patch size | 0.14 | 84 |
| Connectivity : Patch size | 0.13 | 84 |
| Connectivity : Distance to outlet | 0.11 | 84 |
| Connectivity : SH | 0.1 | 84 |
| Connectivity : Distance to outlet : SH | 0.01 | 20 |
| Connectivity : SH : Patch size | <0.01 | 20 |
| Distance outlet : SH : Patch size | <0.01 | 20 |
| Connectivity: Distance to outlet : Patch size | <0.01 | 20 |
| Connectivity : Distance to outlet : SH : Patch size | <0.01 | 1 |

**Figure 3:**
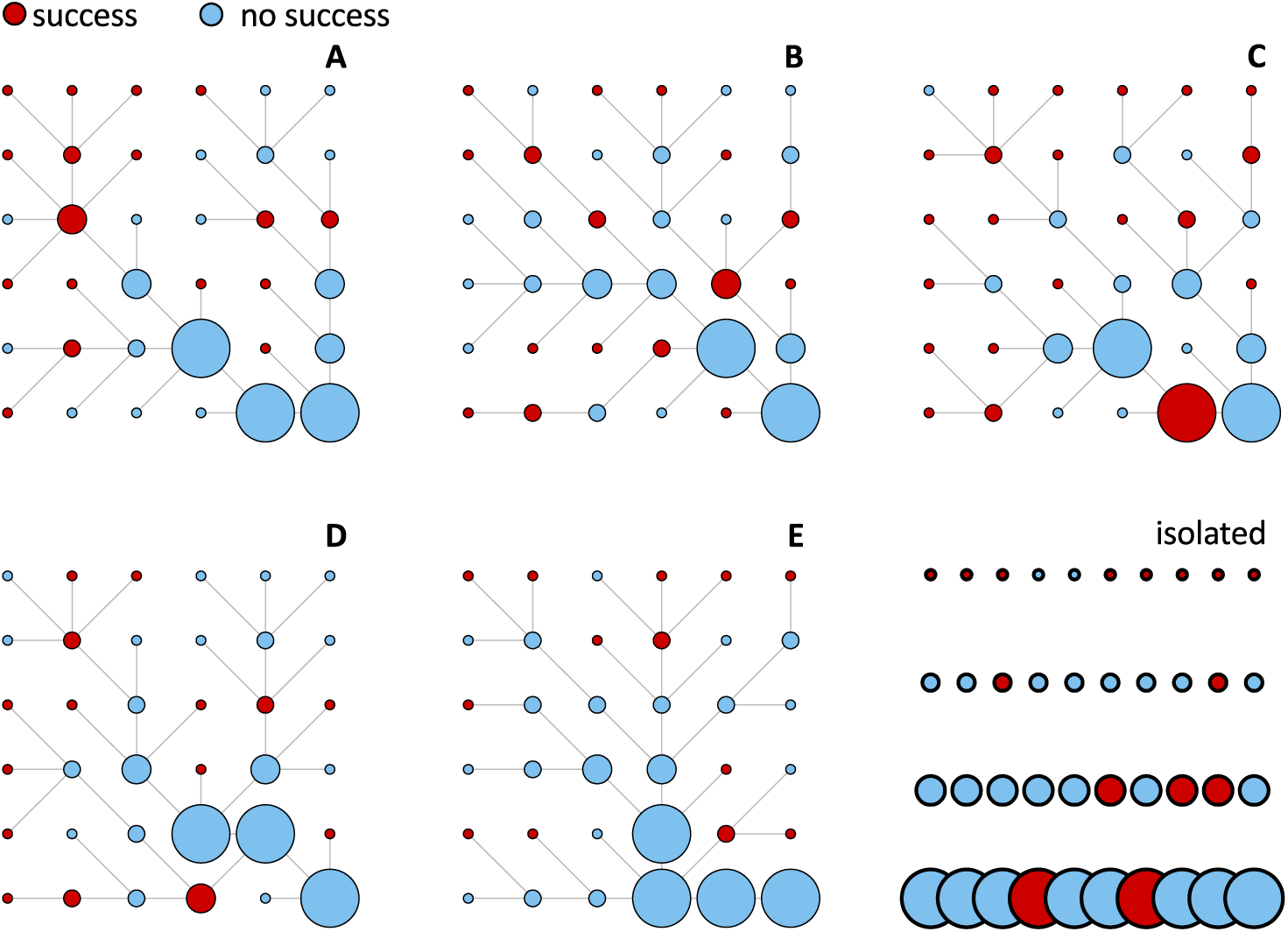
Short-term invasion success of *Spirostomum* sp. across the dendritic landscapes (A to E) and the isolated single patch controls.

Finally, we did model prediction using the best-fitting model for the short-term invasion success including the variables patch size and Shannon diversity (S_H_) immediately preceding invasion to disentangle their relative contribution to invasion success. These predictions support our results, and highlight the relevance of the interaction on patch size distribution and diversity shaped by network position (Fig. 4): the invasion success follows a reverse hump-shaped relationship where invasion success is high at a low S_H_ immediately preceding invasion, then lower at an intermediate S_H_ but finally increases again with high S_H_. Even more, with an increasing patch size gradient, diversity changed invasion success from a negative to a positive correlation.

**Figure 4:**
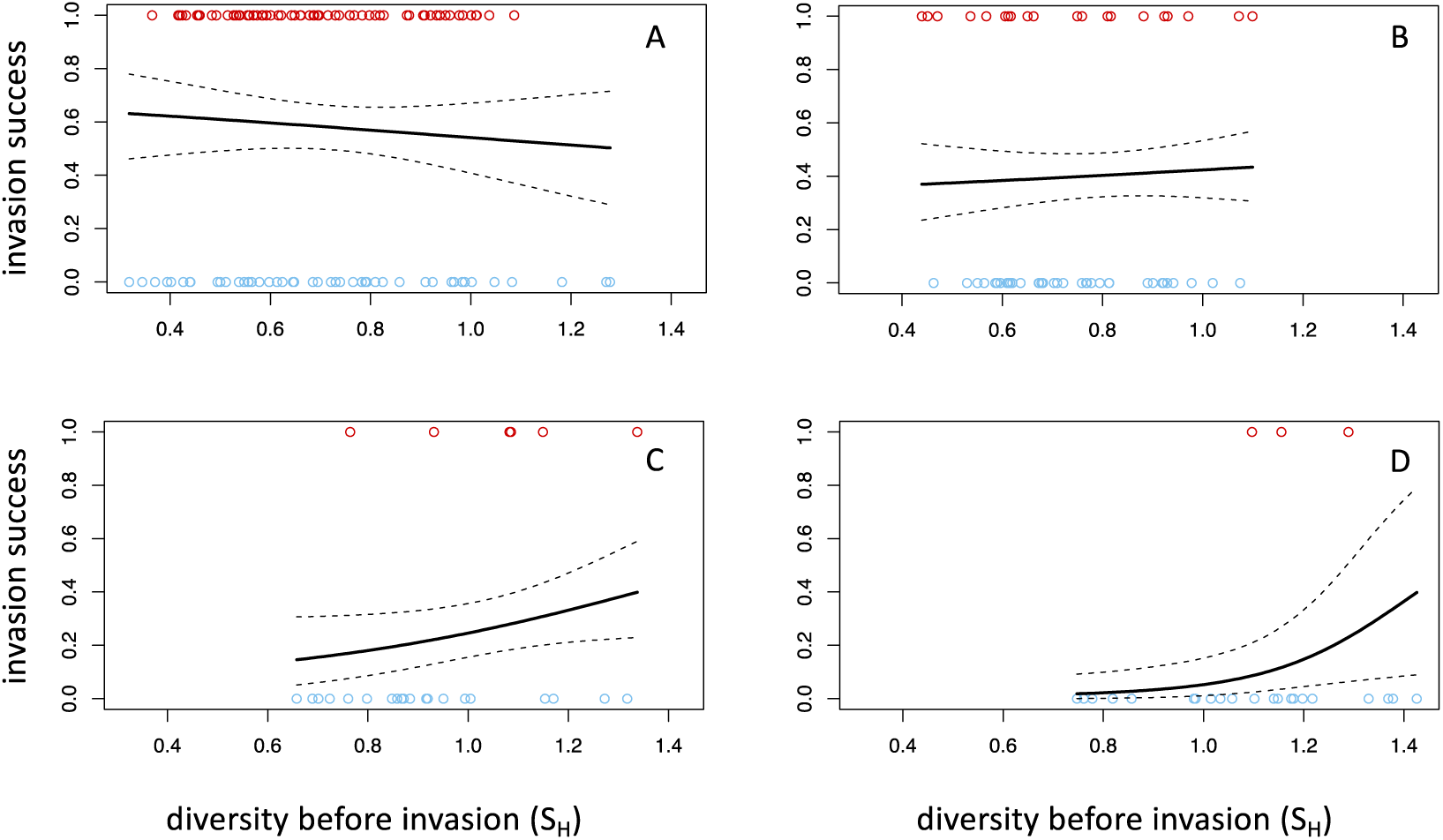
Predictions for short-term invasion success based on the best-fitting glmer model including the variables patch size and Shannon diversity index (S_H_) preceding the invasion across patch sizes (A to D, A = 3 mL, B = 5.2 mL, C = 9 mL, D = 18 mL). Overall, invasion success decreased with increasing patch size (black lines: mean prediction; black dotted lines: 95% confidence interval). However, invasion success was also significantly and non-linear affected by the community diversity preceding the invasion.

## Discussion

Understanding invasion success of species has been a long-standing question in ecology (Elton 1933), and both local properties of the community as well as spatial dynamics/spatial network structures have been used to explain invasion success. We here experimentally showed that these two views are not independent, but can go hand in hand: spatial dynamics, including dispersal and habitat network structure, are shaping local communities, and thereby directly modulating their resistance to invasions. This is likely a strongly understudied, but crucial mechanism: focusing on local or spatial drivers only will not give realistic understanding on how invasions in natural ecosystems emerge.

In our experimental dendritic networks, invasion success was directly modulated by patch size: both short- and long-term invasion success was higher in smaller patches (Fig. 3). However, there was also an indirect effect namely the effect of community diversity (Shannon diversity) preceding the invasion, which was again affected by patch size, but also by the spatial position in the dendritic network: position in the network (as described by patch size, connectivity or distance to outlet) indirectly affected invasion success via an effect on diversity. Thus, communities were shaped and emerged by the interplay of both local dynamics (ecological selection and drift) as well as spatial dynamics (dispersal, in this case along a well-defined and realistic habitat network structure). In short, regional scale properties (here patch size) influence local scale properties (here diversity), which in turn affect the invasibility of local communities to biological invasions.

The direct effect of habitat network structure, and so-called “invasion hubs” has been recently shown in studies of ecological invasions (Morel-Journel et al. 2019). Also, recent experimental and theoretical work indicated that that the spatial percolation of local perturbations will be shaped (and potentially halted) by specific spatial network topologies (Gilarranz and Bascompte 2012, Gilarranz et al. 2015). Our work now demonstrates that this is equally relevant and directly applicable to biological invasions: spatial dynamics shape the local properties, and subsequently affect invasion success. This, as a consequence, will also allow novel strategies to combat and withstand biological invasions. Beyond the direct manual extirpation of the invading species, direct modifications of local conditions, or modifications of the landscape topology to directly halt invasions, we identify a novel strategy: the management of spatial community networks such that spatial dynamics per se modulate local community properties, rendering a better resistance to biological invasions. As such, well connected habitats are shown to be more resistant to biological invasions, likely due to direct effects of connectivity on diversity and structure of local communities, which is a well-known effect in ecology: there are manifold studies showing an increased diversity and more complex local community structure emerging in well-connected landscapes (Damschen et al. 2006, Brudvig et al. 2009, Carrara et al. 2012). This higher diversity then makes the local communities more resistant to invasions. It is thus a “to kill two birds with one stone” situation: appropriate management of spatial networks, and the maintenance of spatial networks *per se*, not only leads to more diverse local communities and maintains diversity as such, but also makes these communities more resistant to biological invasions (Harvey et al. 2016, Bullock et al. 2018).

Importantly, in our experiment we could disentangle the effect of spatial dynamics on shaping local communities and the influence on their invasibility, and the invasion process itself. By definition, the invasion process in natural system has to occur within a spatial context, with invasions spreading through the landscape along specific habitat networks. Importantly, and as a cautionary remark, our study was done in networks in which patch size and connectivity are inherently correlated, and their individual contributions to invasion success cannot be completely teased apart (but see Carrara et al. 2014). We thus specifically highlight the need of simultaneously addressing both components, local and spatial properties, shaping invasion success, as the sole focus on either local dynamics or the spatial unfolding does not cover the herein shown interaction. While invasion success has repeatedly been found to be negatively correlated with community diversity measures (Tilman 1997, Stachowicz et al. 1999, Naeem et al. 2000, Kennedy et al. 2002), there is still some controversy on how this plays out in natural systems, and may be context-dependent.

Biological invasions, and their consequences on natural communities, are likely to become even more important in the future (Ricciardi et al. 2017). By showing that well-known regional scale properties, such as patch size and connectivity distribution in complex landscape networks, as well as local scale properties, such as diversity, affect invasion success, we unite two previously often disconnected lines of argumentation when understanding the (spatial) unfolding of invasions. Most importantly, the intricate effects of spatial dynamics shaping local communities, and making them more or less susceptible to biological invasions highlights that the strategies for prohibiting invasions must go hand in hand: local management of communities as well as maintenance of regional-scale spatial dynamics should be optimized to maintain diverse and complex natural communities, increasing their resistance to biological invasions. Notably, however, the maintenance and management of spatial networks is a two-sided sword: connectivity and spatial dynamics can make local communities more resistant to invasions. However, they are also the way that biological invasions spread in space. While isolation and disconnecting spatial networks may seem a viable strategy to reduce the spread of biological invasions, we believe that this may be short-sighted: the lack of sufficient spatial dynamics will eventually lead to the loss of local diversity, making these communities less resistant to invasions in the long-term. We thus emphasize that it is essential to manage natural ecosystem networks as well as invasive species on a landscape level.

## Author Contributions

All authors planned and designed the study; KH conducted the experiment and collected the data; KH analyzed the data. FA and KH led the writing of the manuscript. All authors contributed critically to the drafts and gave final approval for publication.

## Acknowledgements

We thank Emanuel A. Fronhofer for support during data analysis and Chelsea J. Little for commenting the manuscript. Funding is from the Swiss National Science Foundation Grant No PP00P3_150698 (to F.A.).

