## Appendix for "Patch size distribution and network connectivity affect species invasion dynamics in dendritic networks"

Table S1. Anova on diversity preceding invasion in patches of different sizes and comparing landscapes with controls.

**Df	Sum Sq	Mean Sq	F value	Pr(>F)**

landscape_type	1	0.190	0.190	4.594	0.0332

volume	1	3.365	3.365	81.528	<2e-16

landscape_type:volume	1	0.003	0.003	0.071	0.7906

Residuals	216	8.914	0.041

Figure S1

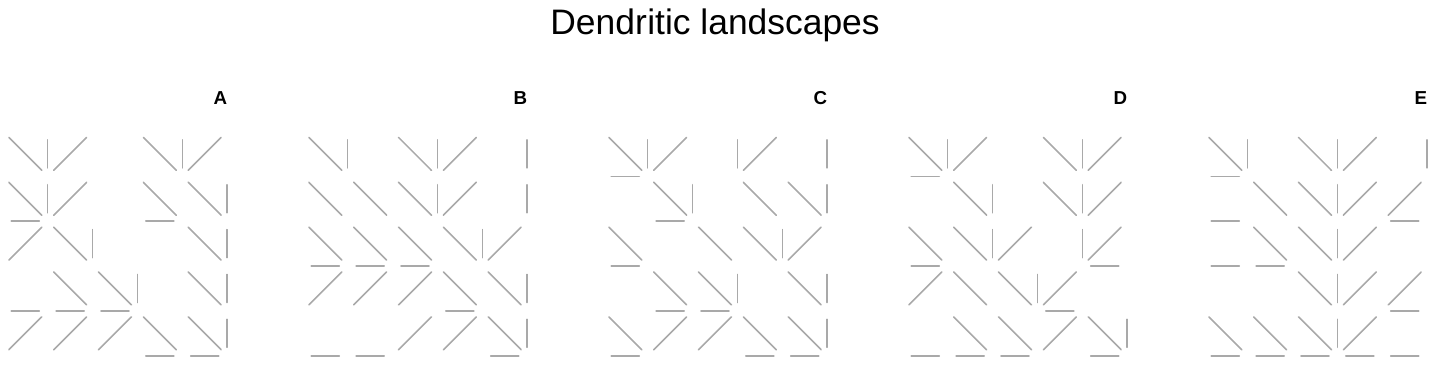

Figure S1: Network structures of the connected dendritic landscapes A to E. Each landscape consists of 36 patches (blue circles). The connection between the patches is indicated by the black lines. Connectivity of the patches was made artificially by pipetting volumes bidirectionally from one to another patch.

Figure S2

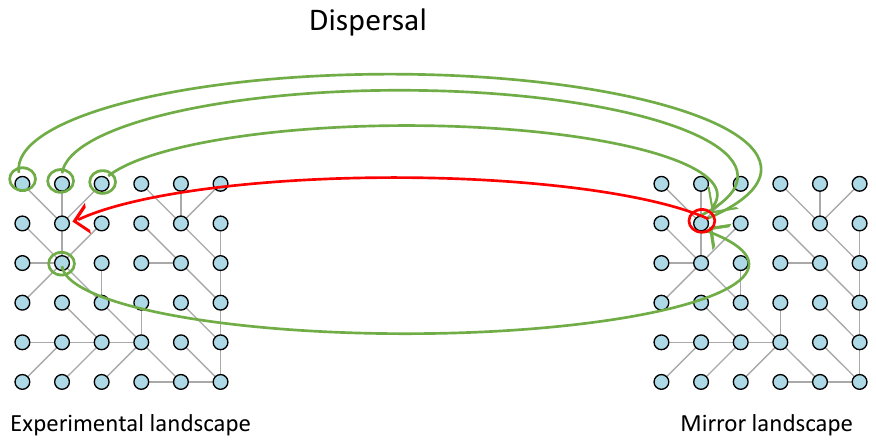

Figure S2: Bidirectional dispersal was conducted along the network structure of the landscapes using a mirror landscape to avoid long-tailed dispersal. Example given for landscape A: From the experimental landscape the volume of 200 µL was taken out each patch (green arrows) connected to the centered one (blue circle) and transferred to a mirror landscape. The mirror landscape followed the same connectivity as the experimental landscape. After well mixing the volumes in the mirror landscape it was transferred back to

Figure S3

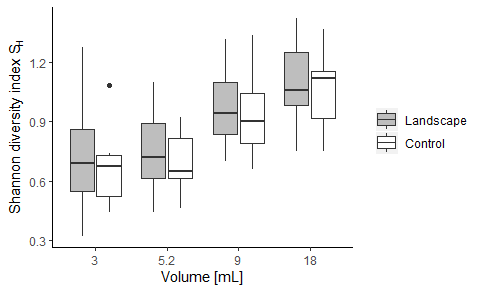

Figure S3: Shannon diversity index (S_H_) before invasion. Diversity is lowest in the small (3 mL) patch (mean = 0.70) and highest in the large (18 mL) patch (mean = 1.07).

Figure S4

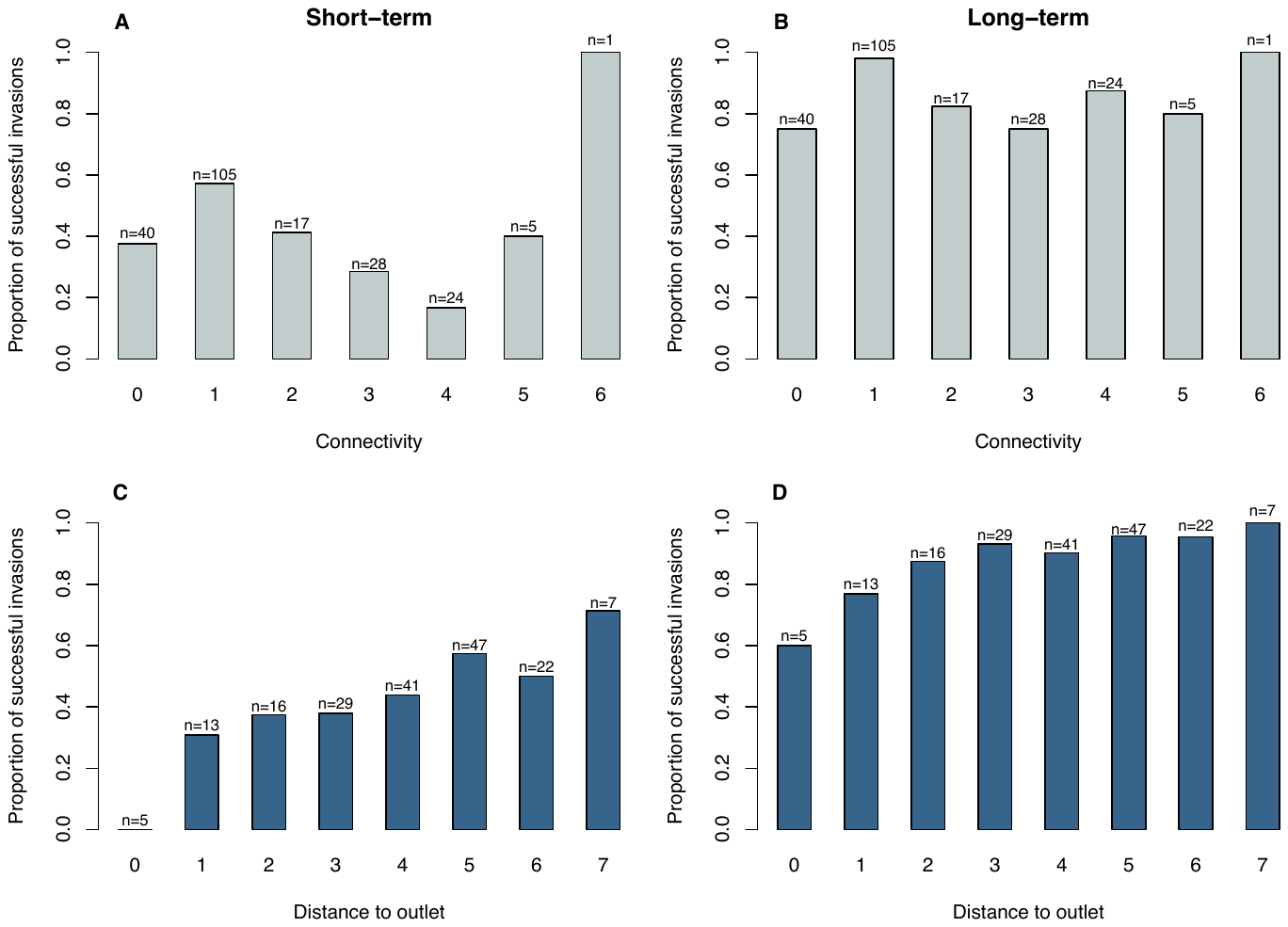

Figure S4: Proportion of invasion success as a function of connectivity. Four days after the invasion of *Spirostomum* sp., a volume of 175 µL was screened for the invader under a stereomicroscope. The isolated single patch control is visualized as connectivity 0, connectivity 1 to 6 are the patches of the landscapes. Numbers above the bars are according to the total numbers of patches an invasion is possible.

Figure S5

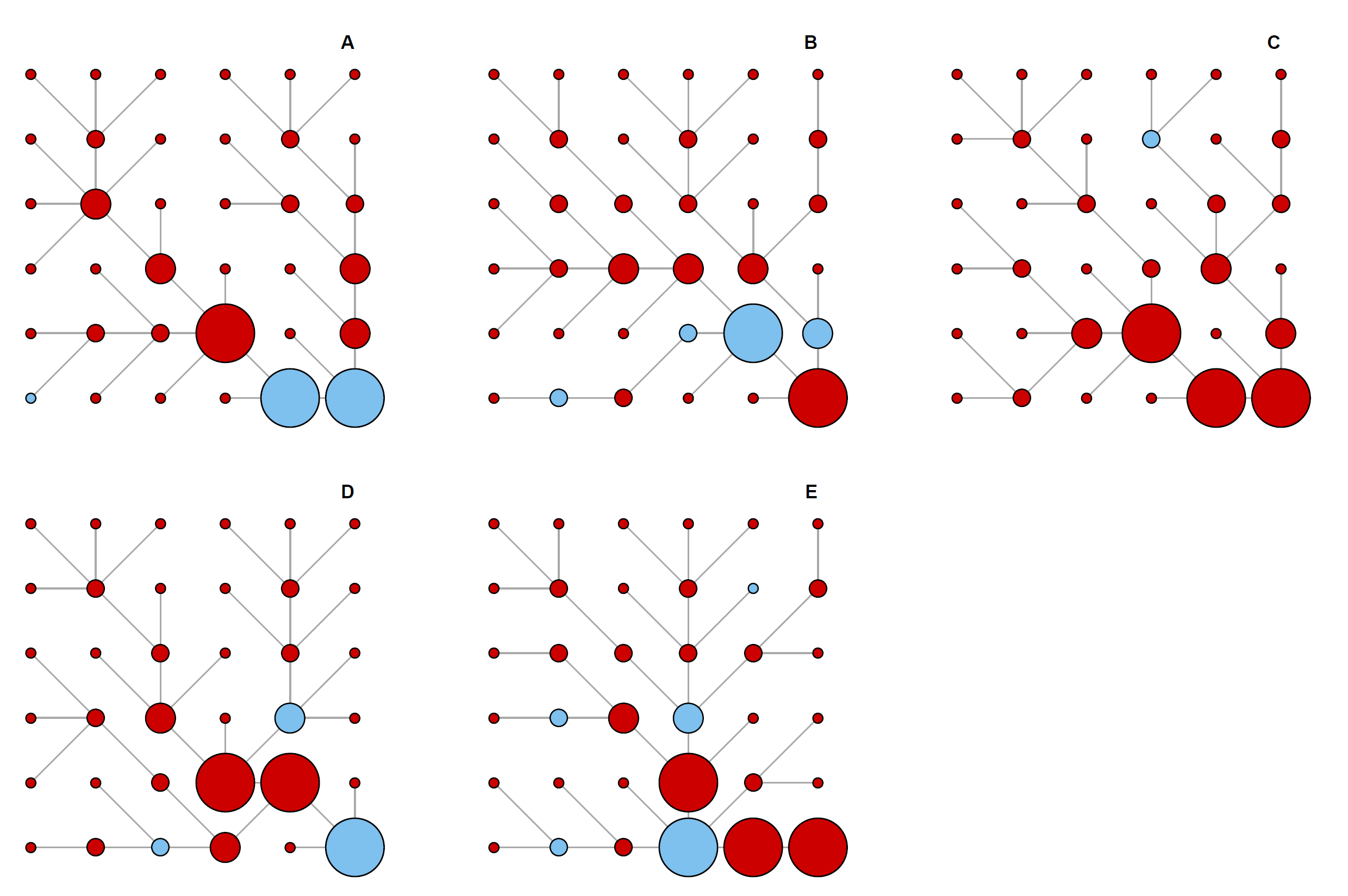

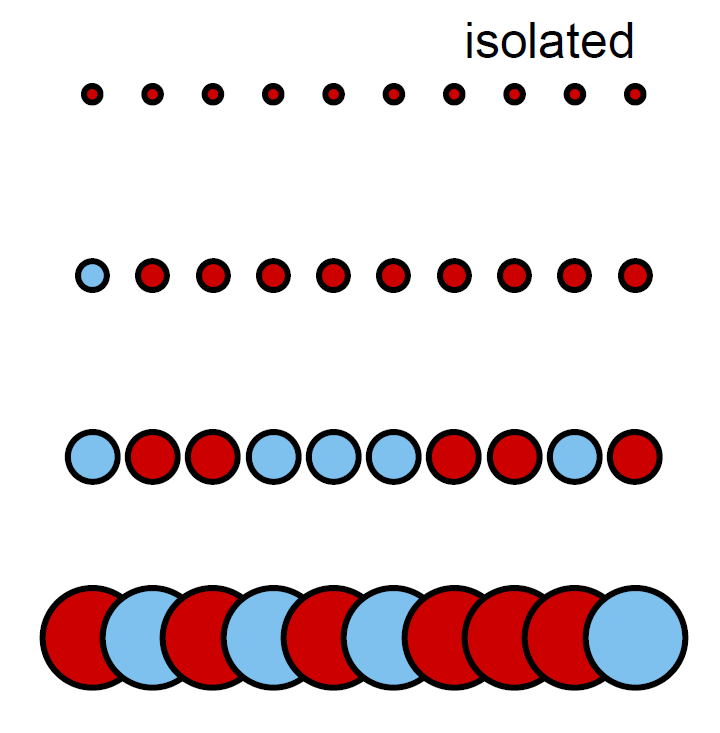

Figure S5: Long-term invasion success of *Spirostomum* sp. across the dendritic landscapes (A to E) and the isolated single patch controls.

Figure S6

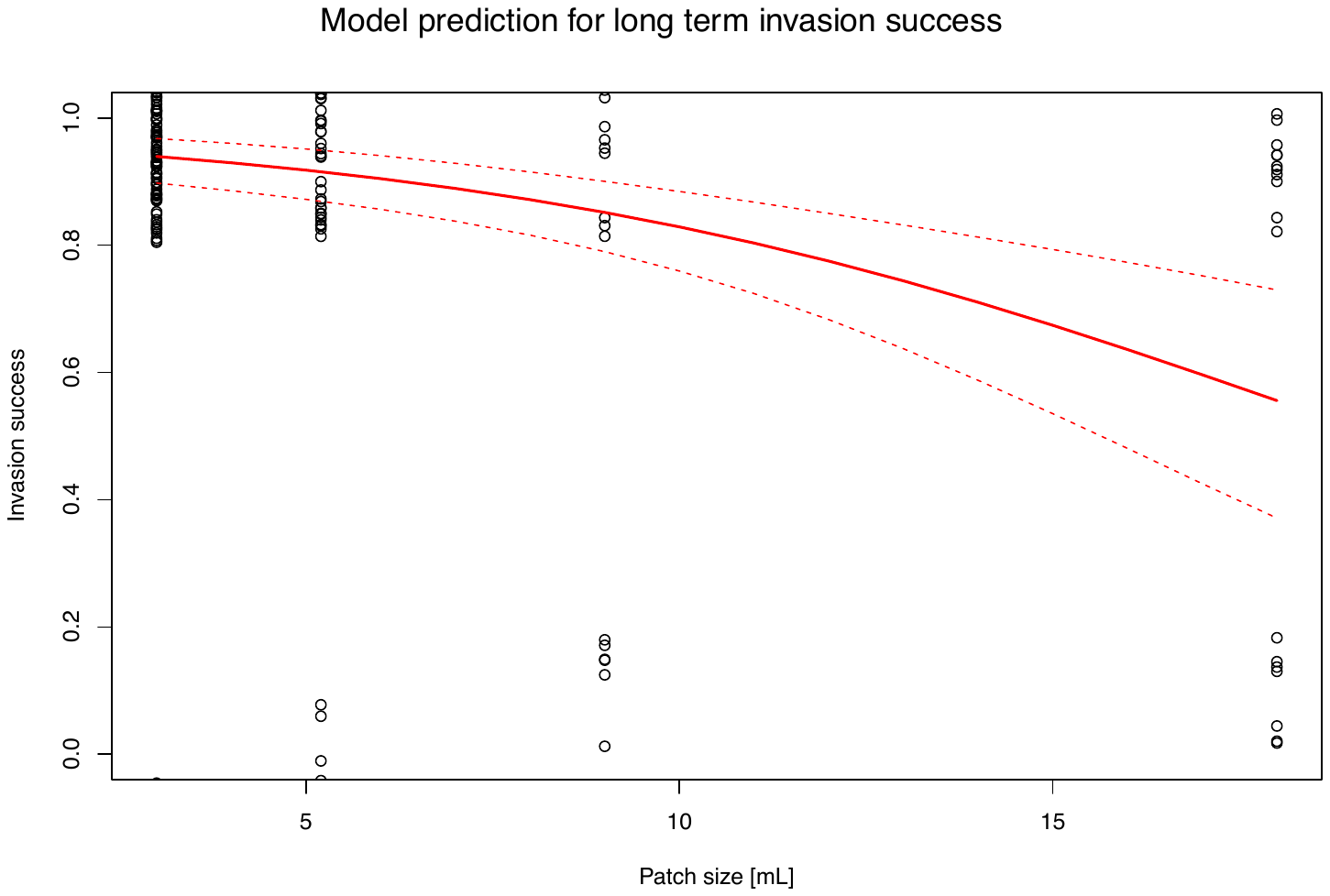

Figure S6: Predictions for long-term invasion success: with increasing patch size the invasion success is predicted to decrease (red line), 95% confidence interval (red dashed line).
